# Sex- and ketogenesis-dependent effects of intermittent fasting against diet-induced obesity and fatty liver disease

**DOI:** 10.1101/2025.11.17.688915

**Authors:** Termeh Aslani, Shaza Asif, Yena Oh, Hyejin Lee, Cole Stocker, Sora Kwon, Julie Pan, Saif Dababneh, Xiaoling Zhao, Letitia Taylor, Adam E. Mullick, Glen F. Tibbits, Ri Youn Kim, Morgan D. Fullerton, Erin E. Mulvihill, Joe Eun Son, Kyoung-Han Kim

## Abstract

Intermittent fasting (IF) improves metabolic health, yet the requirement for hepatic ketogenesis in mediating these benefits remains unclear. Here, we investigated how hepatic ketogenesis contributes to the metabolic and hepatic effects of IF in male and female mice. In the human liver, ketogenesis-associated genes showed sex-dependent correlations with inflammatory and fibrotic pathways. In mice, fasting increased circulating ketone bodies, with females exhibiting a greater rise, indicating intrinsic sex differences in ketone metabolism. IF reduced body weight and adiposity in both sexes, and these systemic benefits persisted despite antisense oligonucleotide (ASO)-mediated knockdown of hepatic *Hmgcs2*. In contrast, hepatic benefits were sex- and ketogenesis-dependent. IF markedly reduced steatosis and fibrosis in male mice, but these improvements were attenuated or abolished when hepatic ketogenesis was disrupted.

Female mice showed minimal hepatic benefit from IF and displayed heightened susceptibility to steatosis, fibrosis, and inflammatory activation under ketogenic insufficiency. Single-cell transcriptomic analyses identified neutrophils and myofibroblasts as key responders to hepatocyte-derived ketone bodies, and IF suppressed neutrophil-driven inflammatory signaling in a ketogenesis-dependent manner in males but not females. Together, these findings demonstrate that while systemic metabolic improvements from IF are largely ketogenesis-independent, the hepatic anti-steatotic and anti-fibrotic effects of IF are sexually dimorphic and require intact hepatic ketogenesis.

## INTRODUCTION

Intermittent fasting (IF) is a well-established dietary intervention that confers health benefits against a range of pathological conditions in humans and preclinical models (1–4). In particular, IF has been shown to effectively prevent and attenuate obesity and associated metabolic dysfunction through reductions in body weight and fat mass, promotion of adipose thermogenesis, and anti-inflammatory remodelling (4–7). Nevertheless, it remains elusive whether IF provides universal benefits across different age groups and sexes. Specifically, although biological sex is a significant determinant of metabolic regulation (8–10), and a recent clinical trial has reported sex-dependent differences in metabolic responses to short-term time-restricted eating (11), most preclinical studies of dietary interventions, including IF and ketogenic diets, have been conducted in male rodents (12–14). This highlights the necessity of investigating the sex-dependent effects of IF.

Based on variations in fasting and feeding durations, diverse forms of IF regimens have been utilized and investigated, including alternate-day fasting and time-restricted feeding (1-3; 15). We previously introduced a 2:1 IF regimen, alternating one day of fasting followed by two consecutive days of *ad libitum* (AL) feeding (7; 16; 17). As this provides the fasted mice with sufficient time to compensate for the loss of energy intake, this regimen enables examination of the effects of isocaloric IF independent of caloric restriction. Using this model, we have demonstrated that 2:1 IF prevents high-fat diet (HFD)-induced obesity and associated metabolic dysfunction in male mice through adipose thermogenesis and improvements in glucose homeostasis (7; 16; 17). 2:1 IF also reduces hepatic lipid accumulation induced by HFD (7), yet the underlying mechanism by which IF confers protection against metabolic dysfunction-associated steatotic liver disease (MASLD) remains poorly understood.

Activated by food deprivation, ketone body metabolism constitutes one of the common underlying pathways of various dietary interventions (18–20). Fasting and IF can stimulate a state of physiological ketosis through controlled and regulated increases in ketone body production via fatty acid oxidation (i.e., ketogenesis), primarily in the liver, generating β-hydroxybutyrate (β-OHB), acetoacetate, and acetone. These KBs can then be used in other organs as a source of energy. As mitochondrial hydroxy-3-methylglutaryl-CoA synthase II (Hmgcs2) is the rate-limiting enzyme of ketogenesis, we and others have shown that ketogenic dysfunction via *Hmgcs2* ablation leads to spontaneous fatty liver disease in postnatal mice, whereas ketogenesis activation via HMGCS2 overexpression mitigates HFD-induced hepatosteatosis (21–25). These findings suggest a causal role of hepatic ketogenesis in fatty liver disease development, progression, and treatment; however, it has not been tested whether ketogenesis is required for IF-mediated metabolic benefits, particularly against MASLD.

In this study, we tested whether 20 weeks of 2:1 IF could protect against obesity and fatty liver disease in HFD-fed male and female C57BL/6J mice lacking hepatic ketogenesis through antisense oligonucleotide (ASO)-based *Hmgcs2* knockdown. We found that ketogenesis insufficiency did not impair the anti-obesity effect of IF in either sex. Notably, hepatic Hmgcs2 knockdown attenuated the anti-steatotic effects of IF in male mice, whereas in females, loss of ketogenesis largely abolished IF-mediated protection against steatosis. More importantly, the anti-fibrotic effects of IF were abolished in ketogenesis-deficient mice of both sexes.

Bioinformatic analysis using a mouse liver single-cell RNA-sequencing (scRNA-seq) dataset, coupled with experimental validation, indicated that ketone body metabolism was linked to inflammatory signalling pathways, thereby modulating hepatic fibrosis. Aanalyses of human liver transcriptomic data further supported the translational relevance of the ketogenesis-inflammation-fibrosis axis. Together, these findings demonstrate that IF confers metabolic protection in sex- and ketogenesis-dependent manners, providing mechanistic insights into this widely adopted dietary intervention

## MATERIALS AND METHODS

### Animal maintenance

All animal experiments were conducted in accordance with protocols (#2950, #4013, and #2962) approved by the Animal Care Committee in the Animal Care and Veterinary Service at the University of Ottawa and conformed to the guidelines of the Canadian Council on Animal Care. All mice were housed in standard ventilated cages in temperature- and humidity-controlled rooms with 12-hour light-dark cycles (21-22 °C, 30-60% humidity for housing), and free access to water. The C57BL/6J mice used in this study were generated from breeders obtained from Jackson Laboratory (#000664). After weaning (3-week-old) to 8-week-old, mice were fed standard chow in which 22% of calories were from fat, 55% from carbohydrates and 23% from protein (2019 Teklad Global Diet; Envigo). From 8-week-old, mice were fed a high-fat diet (HFD) in which 45% of calories were from fat, 35% from carbohydrates, and 20% from protein (D12451; Research Diets).

### *Hmgcs2* knockdown in mice

To ablate fasting-induced ketogenesis in mice, we used antisense oligonucleotide (ASO)-mediated knockdown of the mouse *Hmgcs2* gene, as previously reported (22; 26; 27).

Specifically, 8-week-old C57BL/6J mice were injected intraperitoneally (i.p.) with 25 mg/kg of murine *Hmgcs2*-targeted ASO (5’-CTGTTTGTCACTGCTGGATG-3’) every 12 days (specifically at the fasting cycle of the IF regimen). The control ASO (141923; 5’-CCTTCCCTGAAGGTTCCTCC-3’), which has no complementarity with any known genes, was employed to demonstrate the specificity of target reduction.

### 2:1 intermittent fasting

The 2:1 IF regimen consists of one day of fasting followed by two days of normal eating (7; 16; 17). Mice were evenly divided into 2 groups: AL and IF; or 4 groups: control ASO AL, control ASO IF, Hmgcs2 ASO AL, and Hmgcs2 ASO IF. 2-3 mice were housed in each cage. After 1 week of acclimation to a new feeding environment, the IF regimen commenced with a fasting cycle in which mice were transferred to a clean cage at 12 PM. Food was not added to the IF group, while a weighted chow was provided to the AL group. After 24 hours (12 PM the next day), the body weights of all the mice were measured, and a weighted chow was added to the IF group. After 48 hours of the feeding cycle, body weights and the amount of remaining chow were measured in both AL and IF groups. This 2:1 IF cycle was repeated throughout the course of the study: 3 weeks for the short-term study and 20 weeks for the long-term study.

### Metabolic phenotyping

The body composition of mice was measured using the EchoMRI-3-in-1 machine (Echo Medical Systems, Houston, TX, USA), as previously described (16). Blood ketone levels were measured using a ketone meter (Freestyle Optium Neo and β-ketone Test Strips; Abbott Diabetes Care Ltd.). Liver triglyceride measurement was performed using the Triglyceride Colorimetric Assay Kit (Cat# 10010303; Cayman Chemical). Briefly, frozen liver tissues (25 - 35 mg) were homogenized in 500 μL of 1× NP-40 Substitute Assay Reagent using a Mini-Beadbeater (BioSpec Products). After 10 min of centrifugation at 10,000 × g at 4°C, the supernatants were diluted 1:5 in the NP-40 Substitute Assay Reagent. The triglyceride content in the solubilized samples was quantified by measuring the absorbance at 540 nm.

### Histological analysis of adipose and liver tissues

Adipose tissues, including inguinal white adipose tissue (IWAT), perigonadal white adipose tissue (PWAT), brown adipose tissue (BAT), as well as liver tissues, were fixed in 4% paraformaldehyde, embedded in paraffin, and sectioned at 5 μm thickness. Tissues were stained with hematoxylin and eosin (H&E) or picrosirius red. Slide images were captured using the Aperio VERSA 8 Scanner (Leica Biosystems).

Areas of picrosirius red-stained fibrosis sites were quantified using Image J software (28). The percentage of fibrosis area was determined by comparing the collagen-stained area to the total tissue area within each field. Images were acquired at 20× magnification, converted to grayscale, and thresholded to identify the collagen-positive area. For each liver, four non-overlapping fields were analyzed, and the mean value was used to represent the percentage of fibrosis area. Adipocyte areas were quantified using ’Adiposoft (v.1.16)’ ImageJ (v.154f) plugin (29) on three representative images at 20× magnification per sample. After manually adjusting the black/white color curve to ensure consistency, images were automatically analyzed, employing 0.5 µm/pixel scale and adipocyte diameter thresholds ranging from 19 µm to 100,000 µm.

### RNA isolation and quantitative RT-qPCR

Frozen tissue samples in TRIzol^TM^ Reagent (Invitrogen) were homogenized using a Mini-Beadbeater (BioSpec Products). RNA was extracted and purified using the PureLink^TM^ RNA Mini Kit (Invitrogen). 2 μg of RNA was reverse-transcribed into complementary DNA (cDNA) using the High-Capacity cDNA Reverse Transcription Kit and RNaseOUT^TM^ (Invitrogen). Gene expression assay was conducted with 5 ng/μL cDNA using Power SYBR Green Master Mix (Thermofisher) on Quant Studio 5 (Applied Biosystems). Relative cycle threshold (CT) values were normalized to the housekeeping TATA-Binding Protein (*Tbp*) gene. For direct comparison of gene expression levels, all samples were analyzed on a single quantitative PCR (qPCR) plate. All primer sequences are shown in **Supplemental Table 1**.

### Immunoblotting

Liver tissue lysates were prepared using RIPA Lysis and Extraction Buffer (#89900, Thermo ScientificTM) and protease and phosphatase inhibitor cocktail (#78442; Thermo ScientificTM). After protein quantification using the Pierce Rapid Gold BCA Protein Assay Kit (#A53225; Thermo ScientificTM), 10 or 20 μg of protein samples was used for electrophoresis on a 10% SDS-PAGE gel and transferred onto PVDF membranes (#162011, Bio-Rad). Membranes were incubated with anti-Hmgcs2 antibody (1:1000, sc-393256, Santa Cruz Biotechnology) and β-actin (1:5000, ab8226, Abcam). Blots were developed using ClarityTM Western ECL Substrate (#170-5060; Bio-Rad) and imaged with a ChemiDoc™ Imaging System (Bio-Rad). Band intensities were quantified using Image Lab software (Bio-Rad).

### Bioinformatic analysis for single-cell RNA-seq and cell-cell communications

Single-cell data were processed and analyzed using Seurat (v5.3.0). Metadata and raw counts from mouse liver single-cell RNA sequencing data in the HFD-induced MASLD model (GSE166504) were used to create a Seurat object. Chow diet, 15-week HFD, and 30-week HFD groups were used for downstream analysis, while the 34-week HFD group was removed from the analysis due to a lower number of total and specific cells (i.e., stellate cells). Gene expression counts were normalized by log-transformation with NormalizeData, and the top 2,000 variable features were identified using FindVariableFeatures with the vst method. These features were scaled with ScaleData while regressing out ‘food,’ ‘capture,’ and ‘animal’ as annotated in the original study (30). Linear dimensional reduction was performed with RunPCA followed by Uniform Manifold Approximation and Projection (UMAP) with RunUMAP using the top 20 principal components. A nearest-neighbor graph was generated with FindNeighbours using 20 principal components and cell clusters were identified using FindClusters with a resolution of 0.1.

To investigate cell-cell communication inferences based on metabolomic and ligand-receptor interactions, we utilized MEBOCOST (v1.2.2) (31) and CellChat (v2.2.0) (32), respectively. MEBOCOST objects were derived from Seurat objects subsetted into Chow diet and 15-weeks HFD. Subsequently, UMAP embeddings, RNA counts with GetAssayData (Seurat), gene names from counts matrices, and metadata were extracted. Lowly abundant cell types in all three conditions were removed, and the following cell types were used for downstream analysis: ’Monocyte/MDM’, ’Neutrophils’, ’Kupffer cells’, ’Hepatocytes’, ’Stellate cells’, ’Myofibroblasts’. Separate MEBOCOST objects were generated for each diet condition with the following parameters: cutoff_exp = 1, cutoff_met = 0.2, cutoff_prop = 0.1. Parameters were tested through a grid search, optimizing for the visualization of a dynamic range of metabolite communication networks, across all diet conditions. Parameters selected filtered out cells with less than 20% abundance of a particular metabolite, and groups of cells, in which less than 10% of the cells expressed a particular sensor, or metabolite. Communication events were inferred without further filtering, by shuffling cell labels over 1000 iterations, to determine *p*-values by false discovery rate - permutation test. Significant (a = 0.05) normalized metabolite communication scores were visualized by MEBOCOST FlowPlot. Line width, node size, and *p*-value scales were manually defined, enabling comparison of communication scores between diet conditions.

CellChat objects were generated from Seurat objects of liver samples under chow diet and 15-week of HFD using createCellChat. The CellChatDB.mouse database, excluding ‘non-protein signaling’ pathways was used. SubsetData was performed to only retain genes involved in the database pathways to save computational cost. Following the developer’s protocols (32), over-expressed genes and interactions were identified using IdentifyOverExpressedGenes and IdentifyOverExpressedInteractions, respectively. Communication probability was calculated using computeCommunProb using the trimean method. Pathway-level communication probabilities were determined using computeCommunProbPathway and were aggregated using aggregateNet. Network centrality scores were computed with netAnalysis_computeCentrality. Finally, CellChat objects from chow diet and 15 week-HFD liver samples were merged for downstream comparative analysis.

All scripts used to generate the results and figures presented in this study are available on GitHub (https://github.com/hankimlab/2025-Intermittent-fasting). Other codes are available from the authors upon request.

### Genome-wide gene expression correlation analysis in the human liver

RNA-seq data of 226 human liver samples (161 males and 65 females) were obtained from the Genotype-Tissue Expression (GTEx; V8) project portal (https://gtexportal.org/home/) on 12/01/2023. Genes that are not expressed in any of the liver samples (0 TPM) were excluded. For genome-wide correlation analysis, Pearson’s linear correlation constants (ρ) and *P*-values between *HMGCS2* and all genes were computed using MATLAB (version 2016a). Heatmap was generated by the ComplexHeatmap R package using samples (rows) ordered based on *HMGCS2* expression and genes (columns) ordered based on correlation constants, as previously described (7). For each gene, TPM values were standardized. Pathway enrichment analysis was conducted using positively (ρ > 0.3) and negatively (ρ < -0.25) correlated genes with statistical significance (*P* < 0.05) using the Database for Annotation, Visualization and Integrated Discovery (DAVID) (33).

### Statistical Analysis

All data were presented as mean ± standard error of the mean (SEM). One-way, two-way, or two-way repeated-measures analysis of variance (ANOVA), followed by Tukey’s *post hoc* analysis for multiple comparisons, was used as appropriate. Statistical comparison of correlations was performed using the Cocor R package (34). *P* values less than 0.05 were considered statistically significant. All statistical analyses were performed with Prism 10.0 software (GraphPad Software) and R Studio.

## RESULTS

### Ketogenic gene *HMGCS2* expression in the human liver negatively correlates with inflammation and fibrosis

The expression level of the rate-limiting ketogenic enzyme, HMGCS2, in the liver is positively associated with hepatic ketogenic activity (21). To define pathways linked to hepatic ketogenesis, we utilized RNA-seq data from 226 human liver samples (161 males and 65 females) from the Genotype-Tissue Expression (GTEx) project (**Fig. 1A**). Since expression levels of *HMGCS2* and other key ketogenic genes (*ACAT1*, *HMGCL*, and *BDH1*) did not differ between males and females (**Fig. 1B**), we performed a co-expression analysis of hepatic *HMGCS2* expression to identify pathways linked to hepatic ketogenesis across all samples. This analysis revealed 2,672 positively (ρ ≥ 0.3) and 2,742 negatively (ρ ≤ -0.25) correlated genes with HMGCS2 expression (**Fig. 1B** and **Supplementary Table 2**). Pathway enrichment analysis showed that HMGCS2-positively correlated genes were involved in fatty acid degradation, peroxisomal metabolism, and mitochondrial function, including canonical oxidative genes such as *EHHADH*, *BDH1*, and *ACOX1* (**Fig. 1C, D**; **Supplementary Fig. 1**). In contrast, HMGCS2-negatively correlated genes were enriched for inflammatory and fibrotic pathways, encompassing Toll-like receptor signaling (*IRAK1*, *IRAK3*, *IRAK4*), monocyte activation (*CD44*, *ITGB1*), and innate immune mediators (*TLR2*, *TLR5*, *IFITM2*, *IFITM3*). Notably, neutrophil markers (*S100A8*, *S100A9*, *VNN2*) and fibrosis-related genes (*TIMP1*, *COL4A1*, *ADAMTS3*) were inversely associated with *HMGCS2* expression, suggesting a ketogenesis-anti-inflammation-anti-fibrosis axis in the human liver.

**Figure 1.**
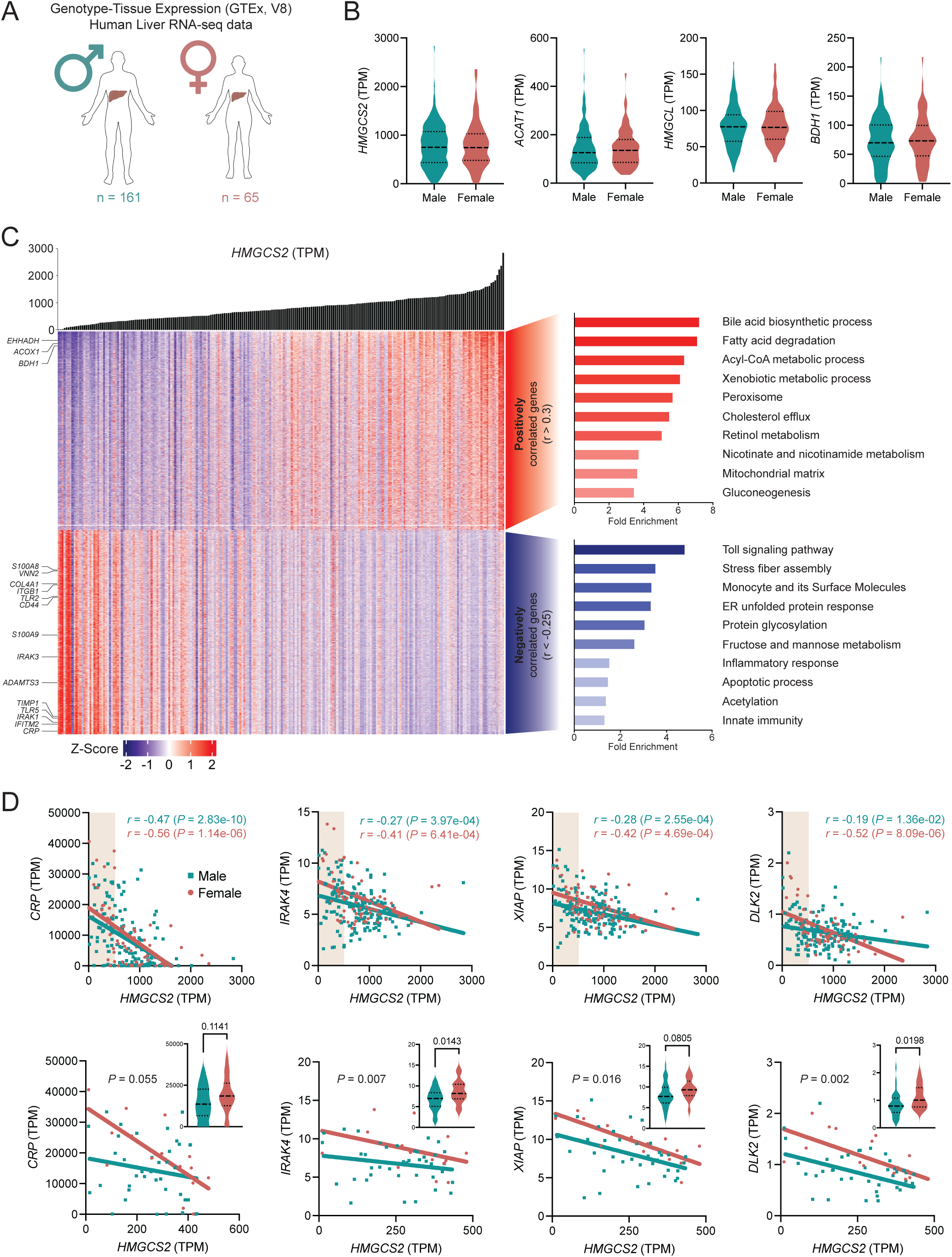
Hepatic HMGCS2 expression in humans associates with metabolic and inflammatory gene programs in a sex-dependent manner. **(A)** Overview of human liver RNA-seq data from the Genotype-Tissue Expression (GTEx, v8) cohort (161 males and 65 females). **(B)** Violin plots showing transcript levels of ketogenic genes (*HMGCS2*, *ACAT1*, *HMGCL*, *BDH1*) in male and female livers. **(C)** Heatmap of genes positively (r > 0.3) and negatively (r < -0.25) correlated with hepatic *HMGCS2* expression levels. Adjacent bar graphs show the enriched pathways among *HMGCS2*-positive (red) and *HMGCS2*-negative (blue) associated genes. **(D)** Representative scatter plots showing inverse correlations between *HMGCS2* and inflammation- or fibrosis-associated genes (*CRP*, *IRAK4*, *XIAP*, *DLK2*) across males (blue) and females (red). The bottom panels show corresponding zoomed-in views to visualize sample-level distribution and sex-specific trends.

We further examined whether these relationships differ by sex and found that several HMGCS2-negatively correlated genes exhibited sex-divergent correlation coefficients (ρ ≤ -0.4 and |ρₘₐₗₑ - ρfₑₘₐₗₑ| ≥ 0.1). These included *CRP*, *IRAK4*, *XIAP*, and *DLK2*, all implicated in inflammatory signaling, immune regulation, and hepatocellular carcinoma progression (35–37). These findings raise the possibility that reduced hepatic ketogenesis may predispose to inflammation and fibrosis in a sex-dependent manner.

### Sex differences in fasting-induced hepatic ketogenesis in mice

To understand sex-dependent differences and the physiological impact of hepatic ketogenesis, we first examined ketogenic actions in mice under fed and fasted conditions. After 24 hours of fasting, Hmgcs2 protein and mRNA levels, as well as other ketogenic enzymes (*Acat1*, *Hmgcl* and *Bdh1*) were comparable between male and female C57BL/6J mice (**Fig. 2A, B**; **Supplementary Fig. 2A**). Notably, plasma β-hydroxybutyrate (β-OHB) concentrations after 24 hours of fasting were significantly higher in females than in males (**Fig. 2C**), indicating a functional sex difference in ketone body metabolism independent of hepatic enzyme expression. These findings further suggest that the physiological and health effects mediated by ketogenesis may be modulated by sex.

**Figure 2.**
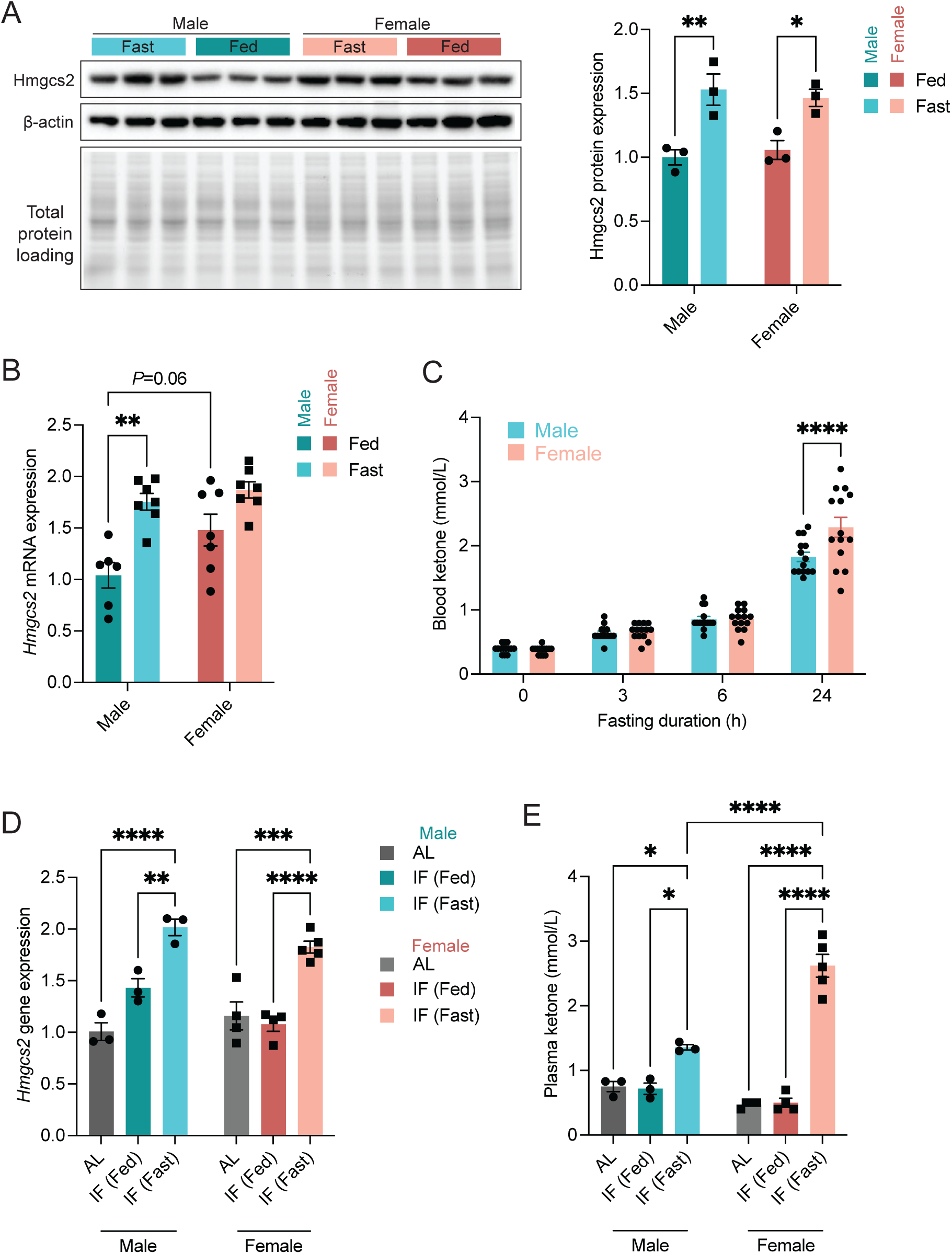
Fasting and IF induce ketogenesis in both sexes, but result in higher circulating ketone levels in females. **(A)** Representative immunoblot (left) and quantification (right) of Hmgcs2 protein expression in male and female mouse livers under fed and fasted conditions. β-actin and total protein loading were used as normalization controls. **(B)** Hepatic *Hmgcs2* mRNA expression after 24 hours of fasting compared to the fed state in male and female mice. **(C)** Plasma ketone body (β-hydroxybutyrate, β-OHB) concentrations in male and female mice after 0, 3, 6, and 24 hours of fasting. **(D)** mRNA expression analysis of the ketogenic gene, *Hmgcs2,* in male and female mice subjected to AL and IF. **(E)** Plasma β-OHB levels measured in groups of AL, IF at feeding and IF at fasting condition in males (n = 3 / group) and females (AL, n = 4; IF (Fed), n = 4; IF (Fasted), n = 5). Statistical analysis was performed by two-way ANOVA or one-way repeated measures ANOVA. *, *P* < 0.05; **, *P* < 0.01; ****, *P* < 0.0001.

While ketone bodies have been considered mediators of the metabolic benefits associated with fasting-mediated dietary interventions, including IF, through various protective mechanisms (18-20; 38), it remains unclear whether hepatic ketogenesis is required for these benefits and whether these effects are sex-dependent. To evaluate early metabolic responses to IF before major changes in body mass and composition, mice were subjected to 2:1 IF regimen for 3 weeks (**Supplementary Fig. 2B**). During this period, hepatic expression of *Hmgcs2* remained similar between sexes (**Fig. 2D**). However, circulating β-OHB concentrations were markedly elevated during the fasting phase relative to *ad libitum* feeding, with females exhibiting a more pronounced increase (**Fig. 2E**). These results further support sex-dependent regulation of ketone body metabolism under fasting and IF.

### Ketogenic insufficiency has no significant effect on IF’s action against diet-induced body weight and adiposity gains in both sexes

We next directly tested whether hepatic ketogenesis is necessary for the metabolic benefits of IF and whether these effects are sex-dependent. Since whole-body *Hmgcs2* knockout (*Hmgcs2*-KO) mice exhibit higher lethargy and mortality between postnatal days 14 and 21 (21), this mouse model is not suitable for adult metabolic studies. To circumvent this limitation, *Hmgcs2* was knocked down using ASO, establishing a ketogenesis-insufficient mouse model (**Supplementary Fig. 3**). 8-week-old C57BL/6J mice were divided into four groups per sex (**Fig. 3A**) and fed a 45% high-fat diet (HFD) under either AL or IF conditions. Control ASO or *Hmgcs2* ASO were administered every fourth IF cycle (*i.e.*, every 12 days), immediately before fasting. Consequently, *Hmgcs2* mRNA and protein expression in the liver were markedly downregulated by *Hmgcs2* ASO in both sexes (**Fig. 3B, C**).

**Figure 3.**
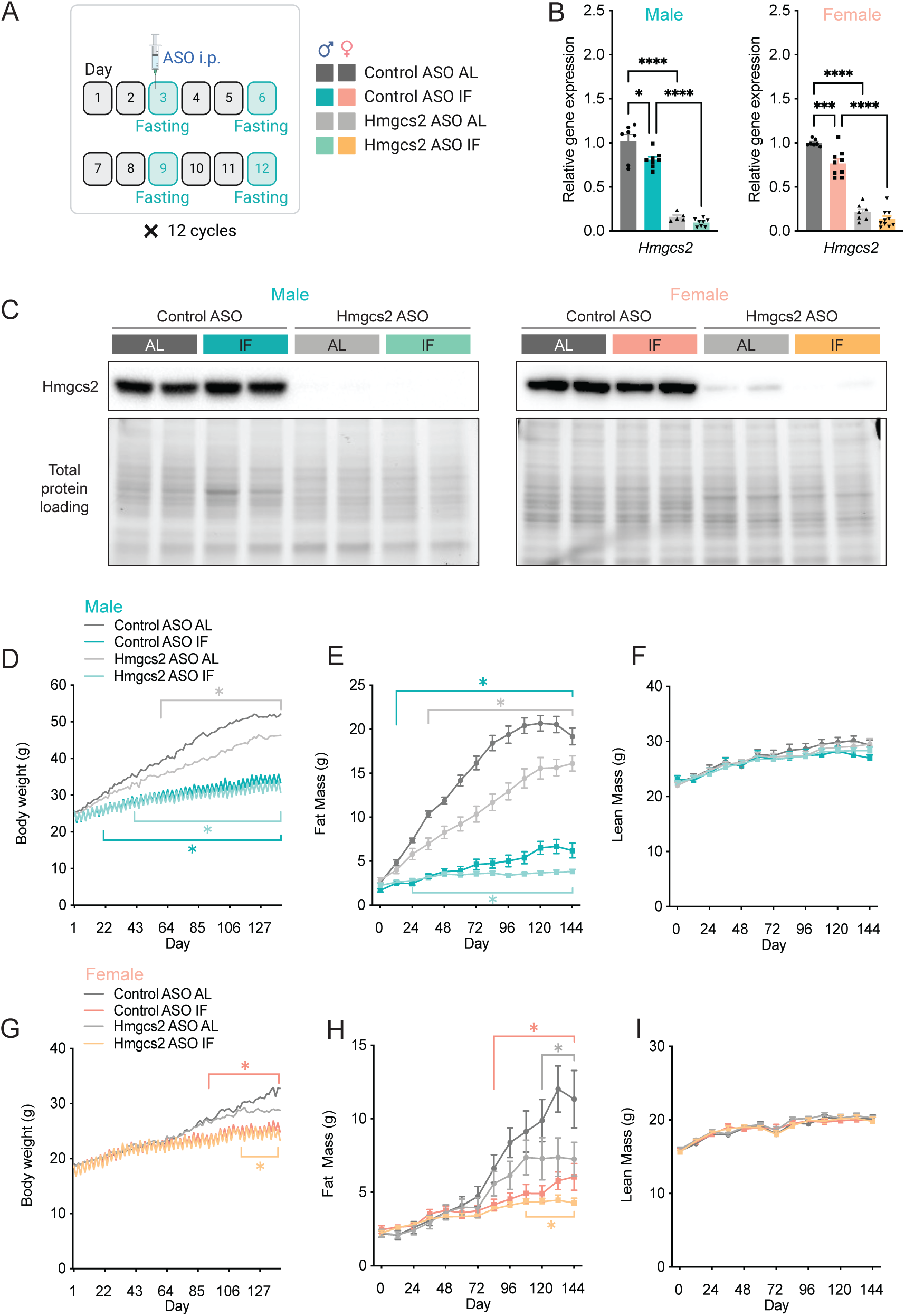
IF attenuates high-fat diet-induced obesity in both male and female mice. **(A)** Schematic illustration of the 2:1 IF regimen with ASO injection. **(B)** *Hmgcs2* mRNA expression in the male (left) and female (right) livers. **(C)** Representative immunoblots showing hepatic Hmgcs2 protein expression and total protein loading controls in male (left) and female (right) mice. **(D-F)** Measurements of body weight (D), fat mass (E), and lean mass (F) in C57BL/6J male mice subjected to IF or AL with control- or Hmgcs2-ASO treatment (Control ASO AL, n = 7; Control ASO IF, n = 8; Hmgcs2 ASO AL, n = 8; Hmgcs2 ASO IF, n = 10). **(G-I)** Measurements of body weight (G), fat mass (H), and lean mass (I) in female mice (Control ASO AL, n = 7; Control ASO IF, n = 9; Hmgcs2 ASO AL, n = 8; Hmgcs2 ASO IF, n = 10). Statistical analysis was performed using a two-way repeated measures ANOVA. *, *P* < 0.05. Green/orange asterisks: Control ASO IF vs. Control ASO AL; light green/light orange asterisks: Hmgcs2 ASO IF vs. Hmgcs2 ASO AL; light grey asterisk: Hmgcs2 ASO AL vs. Control ASO AL. The schematic in Figure 2A was created with BioRender.com.

In control ASO-treated groups, 20 weeks of HFD feeding led to marked increases in body weight (BW) gain (26.5 ± 1.2g) in AL male mice, whereas IF-treated male mice exhibited significantly lower BW gain (10.1 ± 0.8 g) (**Fig. 3D**). Food intake was comparable between control ASO-treated male AL and IF mice (**Supplementary Fig. 4A**), suggesting that IF reduced body weight independent of caloric intake. Body composition analyses showed that lower BW gain in IF-treated male mice was primarily attributed to decreases in fat mass gain (4.8 ± 0.6 g), compared to AL male mice (18.1 ± 1.0 g) (**Fig. 3E**), without changes in lean masses (**Fig. 3F**).

These results are consistent with the previous results seen in C57BL/6J mice (7), indicating no apparent toxicity from control ASO. Compared with control ASO-treated AL males, *Hmgcs2* ASO-treated AL males showed modest but significant reductions in BW gain (20.7 ± 1.1g) and fat mass gain (12.7 ± 1.1g) (**Fig. 3D-F**), consistent with previous findings (22; 26; 27). Notably, in *Hmgcs2* ASO-treated males, IF still markedly decreased HFD-induced BW (8.5 ± 0.5 g) and fat mass (1.4 ± 0.4 g) gains, comparable to control ASO-treated mice, demonstrating that IF retains its anti-obese effects in males even when ketogenesis is impaired.

In females, 20 weeks of HFD feeding induced BW gain (12.9 ± 1.9 g) in control ASO-treated AL mice (**Fig. 3G**). This relative BW gain (67.6 ± 9.6%) were significantly lower (*P* = 0.0002) than in males (105.1 ± 6.5%), consistent with the previous note that female mice are more resistant to diet-induced obesity (39; 40). IF treatment in control ASO females also decreased BW gain (8.2 ± 0.7g; *P* = 0.027), without affecting food intake (**Fig. 3G**; **Supplementary Fig. 4B)**. However, reductions in BW and fat mass only became significant after 96 and 84 days of IF, respectively (**Fig. 3G, H**), indicating a delayed anti-obese response in female mice, compared to male mice. No changes in lean mass were observed with IF in the female mice (**Fig. 3I**). In female AL mice, *Hmgcs2* ASO mildly decreased fat mass during the last 24 days of the IF cycle, but did not affect BW or lean mass (**Fig. 3G-I**). Importantly, *Hmgcs2* ASO-treated females still displayed a mild anti-obese effect of IF, indistinguishable from control ASO-treated females.

As IF significantly reduced fat mass gain in both males and females, we next examined whether hepatic ketogenic function contributes to the anti-obesity effects of IF at the tissue level. Consistent with our previous results (7; 17), IF markedly reduced fat depot weights in control ASO-treated male and female mice (**Supplementary Fig. 5A, B**). Specifically, IF lowered inguinal white adipose tissue (IWAT) and brown adipose tissue (BAT) mass in males, while in females, IF reduced IWAT and perigonadal white adipose tissue (PWAT), not BAT. Histological examination consistently demonstrated that IF prevented adipocyte hypertrophy in IWAT in both sexes (**Supplementary Fig. 5C, D, E, F**). These results were supported by lower expression levels of the adiposity marker gene, leptin (*Lep*), in IWAT from IF-treated males and females (**Supplementary Fig. 5G, H**). Importantly, IF-mediated reductions in fat pad weights, adipocyte size and *Lep* mRNA level were consistently evident in *Hmgcs2* ASO-treated male mice (**Supplementary Fig. 5A, C, D, G**). Similarly, IF decreased adipose tissue size in *Hmgcs2* ASO-treated female mice, although the differences did not reach statistical significance (**Supplementary Fig. 5B, E, F, H**), possibly due to the baseline adiposity-lowering effect of *Hmgcs2* ASO. Together, these data suggest that ketogenesis is not required for IF’s protective effects against HFD-induced obesity and adipose expansion in either sex.

### Hepatic ketogenesis mediates the anti-steatotic and anti-fibrotic effects of IF in the liver of males but not females

Our recent study, along with others, has demonstrated that ketogenic deficiency or insufficiency promotes fatty liver disease under a fat-enriched nutritional environment, whereas activation of hepatic ketogenesis protects against hepatosteatosis (21; 22; 24). We therefore investigated whether ketogenesis is required for the IF-mediated hepatic benefits.

In control ASO-treated male mice, IF markedly reduced liver weight and liver-to-body weight ratio, compared to AL (**Fig. 4A, B**). *Hmgcs2* ASO treatment lowered liver weight in AL males, while liver weight was not further reduced by IF under ketogenic knockdown. Instead, the liver-to-body weight ratio was significantly higher in *Hmgcs2* ASO-IF males, compared to control ASO-IF males. On the other hand, in control ASO-treated female mice, IF had no effect on liver weights, suggesting sex-dependent metabolic impacts. Following *Hmgcs2* knockdown, female mice exhibited a trend toward increased liver weight, compared to control ASO-treated females (**Fig. 4C**). Additionally, liver-to-body weight ratios were significantly higher in *Hmgcs2* ASO-treated AL and IF females, compared to control ASO-treated AL and IF females, respectively (**Fig. 4D**).

**Figure 4.**
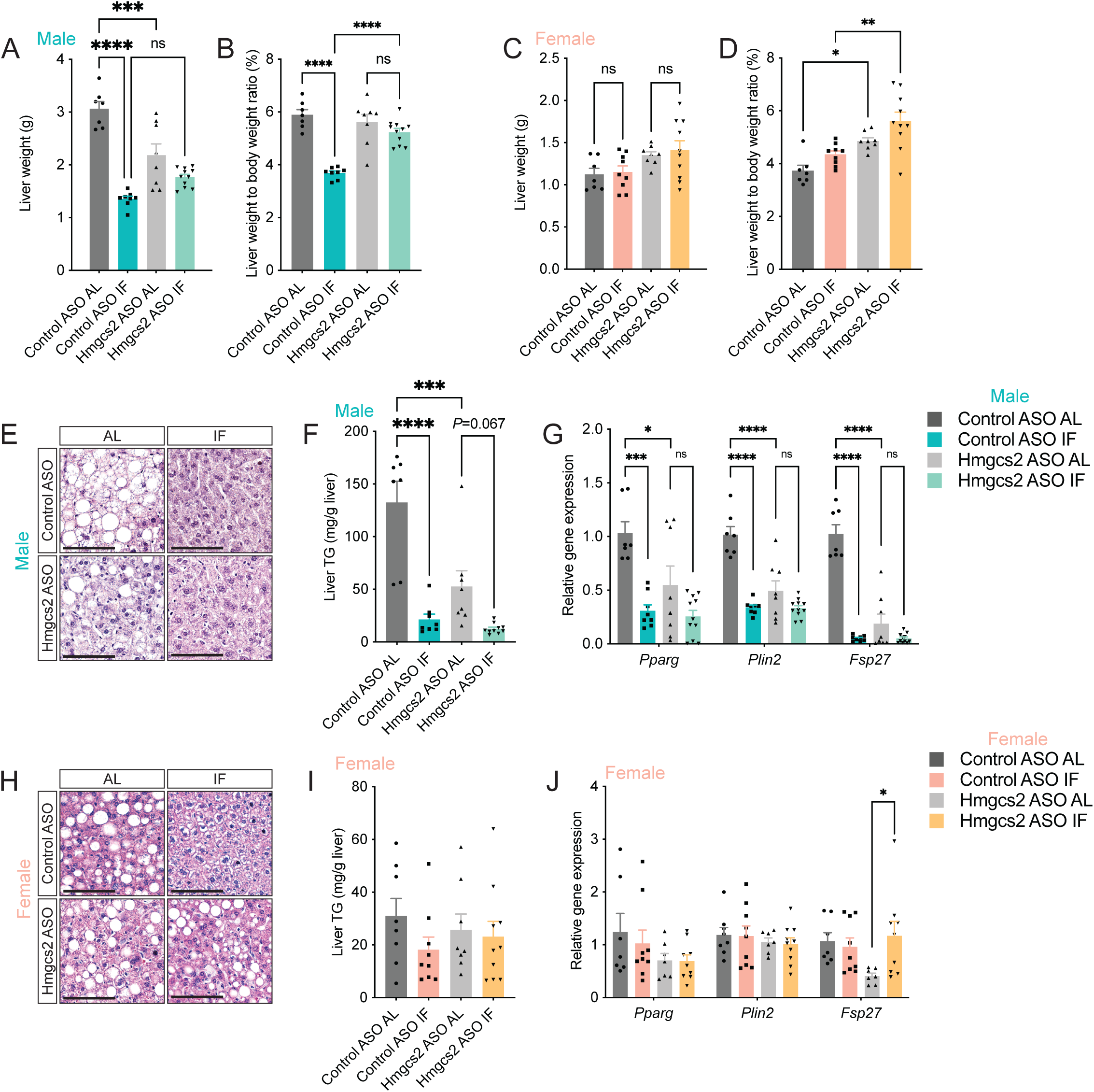
Hepatic ketogenesis drives anti-steatotic benefits of intermittent fasting in male but not female livers. (A-D) Liver weight and liver-to-body weight ratios in male (A-B) and female (C-D) mice subjected to IF or AL with control or Hmgcs2 ASO treatment. **(E)** Representative H&E-stained liver sections from male mice. Scale bar, 100 µm. **(F)** Liver triglyceride content in male mice. **(G)** Relative hepatic expression of lipid accumulation-associated genes (*Pparg*, *Plin2*, *Fsp27*) in male mice. **(H)** Representative H&E-stained liver sections from female mice. Scale bar, 100 µm. **(I)** Liver triglyceride content in the female mice. **(J)** Relative hepatic expression of *Pparg*, *Plin2*, and *Fsp27* in female mice. Statistical analysis was performed by two-way ANOVA. *, *P* < 0.05; **, *P* < 0.01; ***, *P* < 0.001; ****, *P* < 0.0001.

These observations were supported by histological, biochemical, and gene expression analyses. H&E staining and liver triglyceride measurements revealed that HFD-induced hepatosteatosis was markedly reduced by IF in control ASO-treated male mice. However, this protective effect was attenuated and did not reach statistical significance following *Hmgcs2* knockdown (**Fig. 4E, F**). Consistently, IF suppressed the expression of lipid accumulation-associated genes (*Pparg*, *Plin2* and *Fsp27*) in male livers, with a milder effect observed in the *Hmgcs2* ASO group (**Fig. 4G**). In contrast, in female mice, although IF reduced macrovesicular lipid droplets in control ASO-treated livers, microvesicular lipid accumulation within hepatocytes persisted (**Fig. 4H**). This observation was corroborated by unchanged hepatic TG levels and lipid accumulation marker gene expression (**Fig. 4I, J**). Moreover, under *Hmgcs2* knockdown, IF was futile in decreasing lipid accumulation in the female liver. mRNA levels of *Pparg* and *Plin2* were unaffected by IF, while it paradoxically increased *Fsp27*, which regulates lipid storage in the liver (41).

We next assessed whether IF impacts hepatic fibrosis, the primary driver causing liver decompensation in patients with fatty liver disease. Picrosirius red staining showed that IF markedly reduced hepatic fibrosis in control ASO-treated males, but not in *Hmgcs2* ASO-treated males (**Fig. 5A, B**). mRNA levels of fibrosis marker genes, such as *Col1a1* and *Col1a2*, as well as a positive regulator of fibrogenesis *Timp1*, were likewise decreased by IF, while these reductions were attenuated in *Hmgcs2* ASO-treated male liver (**Fig. 5C**). Conversely, IF failed to reduce fibrosis and related marker gene expression in either control or *Hmgcs2* ASO-treated female livers (**Fig. 5D-F**). Notably, *Hmgcs2* knockdown was sufficient to elevate fibrosis genes in both AL and IF female livers. Collectively, these results suggest that in the male mouse liver, IF ameliorates steatosis and fibrosis, with ketogenesis contributing to these protective effects. In contrast, female livers show limited benefit from IF and are more susceptible to ketogenic insufficiency.

**Figure 5.**
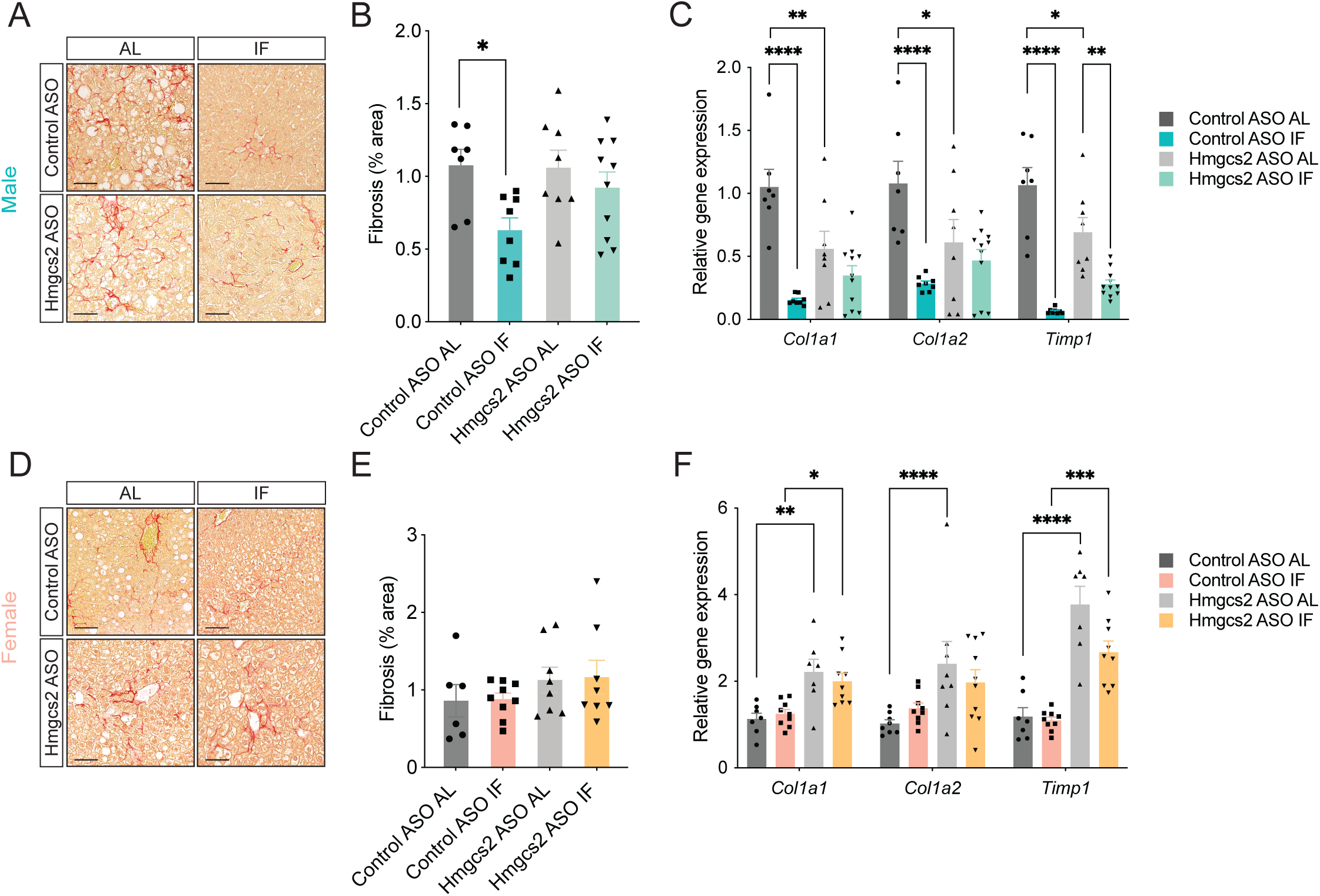
Hepatic ketogenesis drives the anti-fibrotic effects of intermittent fasting in male but not female livers. **(A)** Representative picrosirius red-stained liver sections from male mice treated with control or Hmgcs2 ASO under ad libitum (AL) or intermittent fasting (IF) conditions. Scale bar, 100 µm. **(B)** Quantification of hepatic fibrosis (% area) in male mice. **(C)** Relative hepatic expression of fibrosis-related genes (*Col1a1*, *Col1a2*, *Timp1*) in male mice. **(D)** Representative picrosirius red-stained liver sections from female mice treated as in (A). Scale bar, 60 µm. **(E)** Quantification of hepatic fibrosis (% area) in female mice. **(F)** Relative expression of *Col1a1*, *Col1a2*, and *Timp1* in female livers. Statistical analysis was performed by two-way ANOVA. *, *P* < 0.05; **, *P* < 0.01; ***, *P* < 0.001; ****, *P* < 0.0001.

### Ketone bodies mediate communications from hepatocytes to neutrophils and myofibroblasts in the mouse liver

Hepatic steatosis and fibrosis are driven by complex interactions among various cell types. Prior work has demonstrated that ketone bodies, specifically acetoacetate, inhibit hepatic fibrosis via macrophage-stellate cell communications (26). To further elucidate the cellular and molecular implications of ketone bodies in cell-cell communications during MASLD progression, we analyzed scRNA-seq data from male mouse livers after 15 and 30 weeks of HFD feeding (GSE166504) (30). Clustering analysis identified the major liver cell populations based on canonical marker genes, including hepatocytes (Hep; *Alb*^+^), cholangiocyte/hepatic progenitor cells (HPC; *Epcam*^+^), hepatic stellate cells (HSC; *Reln*^+^, *Rgs5*^+^), myofibroblasts (Myo; *Msln*^+^, *Myl7*^+^), endothelial cells (Endo; *Clec4g*^+^, *Fabp4*^+^), Kupffer cells (KC; *Clec4f*^+^, *Vsig4*^+^), monocytes and monocyte-derived macrophages (MoMF; *Lyz1*^+^, *Ccr2*^+^), neutrophils (Neu; *Retinlg*+, *Ccl4*^+^), T cells (T; *Cd3g*^+^, *Nkg7*^+^), B cells (B; *Cd79a*^+^, *Ms4a1*^+^), natural killer cells (NK; *Xcl1*^+^, *Ncr1*^+^), dendritic cell (DCs; *Flt3*^+^) and plasma dendritic cells (pDC; *Siglech*^+^, *Ccr9*^+^) (**Fig. 6A and Supplementary Fig. 6**). As expected, MASLD progression was reflected by elevated expression of steatotic genes in hepatocytes (*Plin2*, *Cd36*, *Fabp2*, *Apoa4*) and fibrotic genes in stellate cells (*Col1a1*, *Col1a2*, *Timp1*) in 15-week and 30-week HFD livers compared to standard chow-fed control (Chow-fed) liver (**Fig. 6B**).

**Figure 6.**
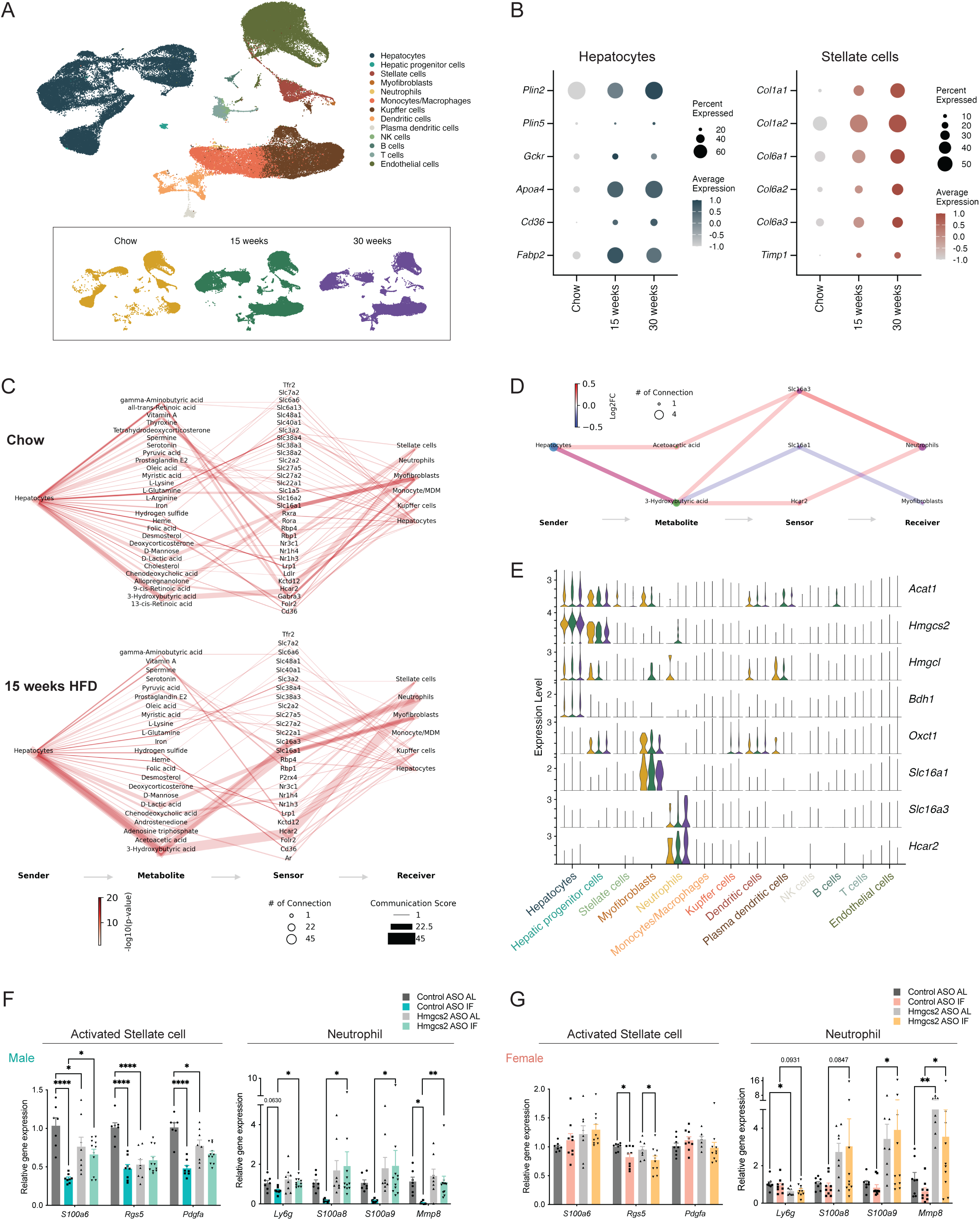
Ketone body-mediated intercellular communication links hepatocytes, stellate cells, and immune cells in the liver. **(A)** Uniform manifold approximation and projection (UMAP) plots of integrated single-cell RNA-seq data from mouse livers under chow diet, 15 weeks HFD, or 30 weeks HFD, annotated by cell type (top) and condition (bottom). **(B)** Dot plots showing expression of representative lipid storage genes (*Plin2*, *Plin5*, *G0s2*, *Cd68*, *Fabp2*) in hepatocytes and fibrosis-related genes (*Col1a1*, *Col1a2*, *Col6a3*, *Col6a2*, *Timp1*) in hepatic stellate cells across dietary conditions. **(C)** Predicted metabolite-mediated cell-cell communication networks derived from hepatocyte-to-non-parenchymal cell interactions under chow and 15-week HFD conditions. **(D)** Simplified network diagram showing representative hepatocyte-derived β-hydroxybutyrate communication pathways through receptors such as *Hcar2* and *Slc16a3*. **(E)** Violin plots showing expression of ketogenic (*Acat1*, *Hmgcs2*, *Hmgcl*, *Bdh1*) and ketone body utilization and transporter genes (*Oxct1*, *Slc16a1*, *Slc16a3*, *Hcar2*) across liver cell populations. **(F, G)** Marker gene expression analysis of activated stellate cells (*S100a6*, *Rgs5*, *Pdgfra*) and neutrophils (*Ly6g*, *S100a8*, *S100a9*, *Mmp9*) in male livers (F) and female livers (G), treated with control or *Hmgcs2* ASO under AL or IF conditions. Statistical analysis was performed by two-way ANOVA. *, *P* < 0.05; **, *P* < 0.01; ****, *P* < 0.0001.

We next examined metabolite-mediated intercellular communication using MEBOCOST (31). The analysis revealed communications via various metabolites from hepatocytes (sender cells) to metabolite-sensors in different receiver cells in Chow-fed and 15-week HFD livers (**Fig. 6C**). Notably, β-OHB and acetoacetate were identified as hepatocyte-derived metabolites targeting two key receiver cell types, neutrophils and myofibroblasts, through ketone body transporters and receptors, such as *Slc16a1* (encoding MCT1, a transporter for ketone uptake in extrahepatic tissues) (42), *Slc16a3* (encoding MCT4, participating in shuttling ketone bodies under fasting or ketogenic states) (43), and *Hcar2* (encoding hydroxycarboxylic acid receptor 2; also known as Gpr109a) (44). Differential cell-cell communication analysis between Chow-fed and 15-week HFD showed that hepatocyte-to-neutrophil ketone body signaling was upregulated, while hepatocyte-to-myofibroblast signaling was downregulated during MASLD progression (**Fig. 6D**). This finding was aligned with the expression patterns of ketone body metabolism genes. Ketogenic genes, such as *Acat1*, *Hmgcs2*, *Hmgcl*, and *Bdh1*, were primarily expressed in hepatocytes, whereas the ketolytic gene, *Oxct1* was enriched in hepatic progenitor cells, myofibroblasts, Kupffer cells, and dendritic cells, but not in hepatocytes (**Fig. 6E**).

Neutrophils expressed *Hcar2* and *Slc16a3*, both increased by HFD, while myofibroblasts expressed *Slc16a1*. Other proposed ketone body transporters/receptors, including *Slc16a6* (MCT7; ketone body secretion in hepatocyte), *Ffar2* (GPR43, short-chain fatty acid receptor for acetoacetate) (38), *Ffar3* (GPR41, receptor for β-OHB) (45) were not significantly detected (data not shown). Together, these results suggest neutrophils and myofibroblasts as primary recipient cells of hepatocyte-derived ketone bodies during MASLD progression.

To determine whether these communications are altered by IF and ketogenic insufficiency, we analyzed neutrophil and myofibroblast markers in our ketogenesis-deficient IF models. In male mice, IF significantly downregulated markers in activated myofibroblasts (*S100a6*, *Rgs5*, *Pdgfa*) as well as neutrophils (*Ly6g*, *S100a8*, *S100a9*, *Mmp8*) under control ASO, but not under *Hmgcs2* ASO knockdown (**Fig. 6F**). In the female liver, IF had little effect, with a mild reduction of *Rgs5* expression (**Fig. 6G**). Notably, neutrophil markers were upregulated by *Hmgcs2* knockdown in both AL and IF female livers, suggesting a heightened susceptibility of immune responses to ketogenic insufficiency. Collectively, these results suggest that in males, hepatocyte-derived ketone bodies mediate IF-induced suppression of myofibroblast activation and neutrophil infiltration, whereas this protective crosstalk was largely diminished in females.

### IF-induced hepatic ketogenesis impacts liver inflammatory cell-cell communications in a sex-dependent manner

During the progression of metabolic dysfunction-associated steatohepatitis (MASH), hepatic fibrosis arises through intercellular communication among hepatocytes, hepatic stellate cells, and diverse immune cell populations (46). Among these, neutrophilic infiltration serves as a critical inflammatory trigger, which activates stellate cells into myofibroblasts and promotes the recruitment and activation of pro-inflammatory macrophages (47). Since our analyses identified neutrophils and myofibroblasts as major cell types responsive to IF-induced ketone bodies, we next examined their interactions with other liver cells during MASLD progression.

To accomplish this, we applied CellChat to the mouse liver scRNA-seq dataset (32), enabling quantitative inference of ligand-receptor interaction networks. Differential interaction analysis revealed that outgoing signals from neutrophils were elevated in 15-weeks HFD livers, whereas signals from myofibroblasts were broadly reduced (**Fig. 7A**). In both cases, Kupffer cells and monocytes/macrophages were the major recipients (receptor expressing cells).

**Figure 7.**
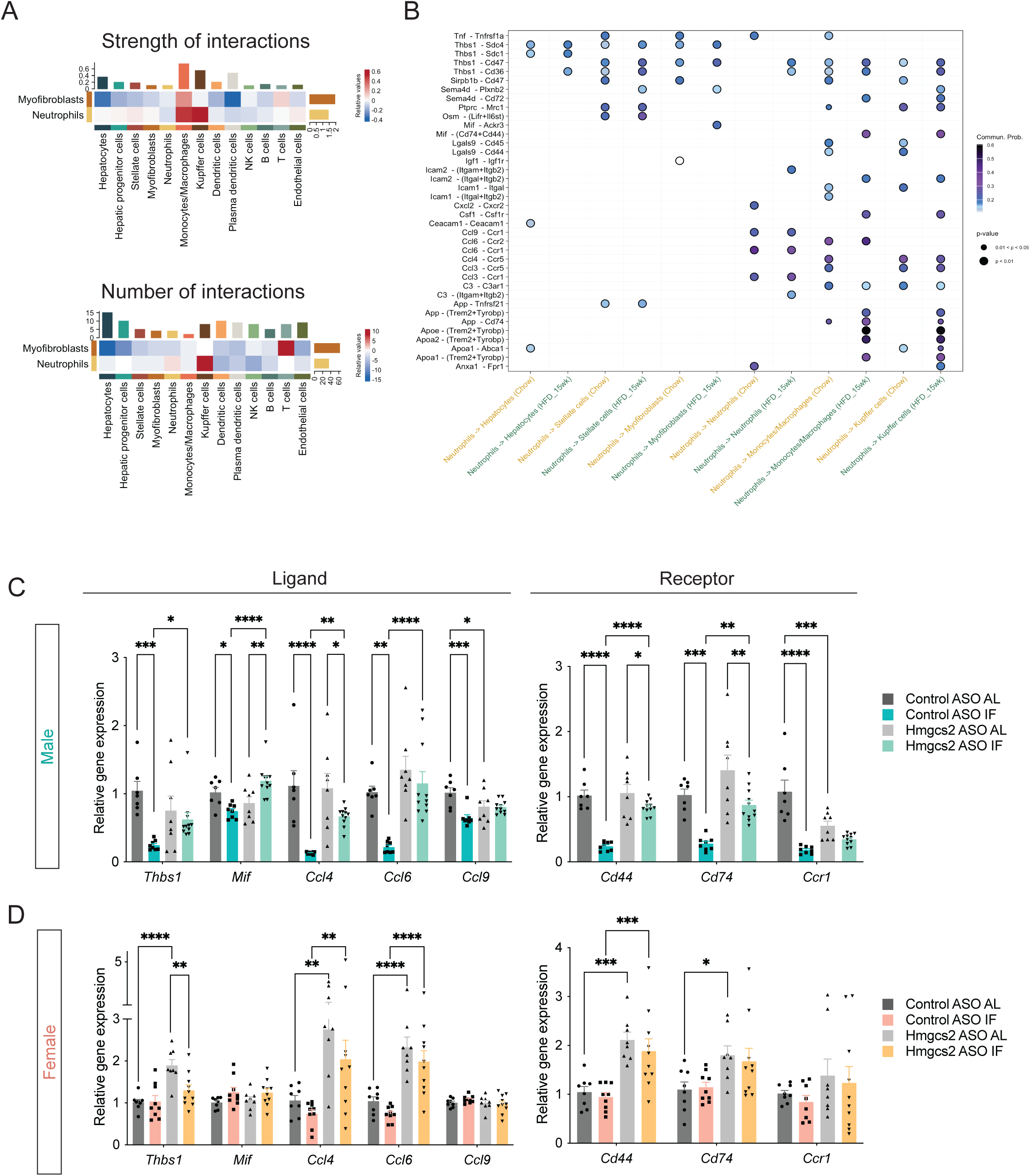
Intermittent fasting suppresses neutrophil-mediated inflammatory signaling and intercellular communication in a sex- and ketogenesis-dependent manner. **(A)** Differential cell-cell interaction analysis between chow and 15-week HFD mouse livers using CellChat. The analysis quantified the strength (top) and number (bottom) of predicted ligand-receptor interactions among major liver cell types. **(B)** Ligand-receptor interactions between neutrophils to other liver cells in chow and 15-week HFD conditions. **(C, D)** Gene expression analysis of inflammatory ligands (*Thbs1*, *Mif*, *Ccl4*, *Ccl6*, *Ccl9*) and receptors (*Cd44*, *Cd74*, *Ccr1*) in male livers (C) and female livers (D), treated with control or *Hmgcs2* ASO under ad libitum (AL) or intermittent fasting (IF) conditions. Statistical analysis was performed by two-way ANOVA. *, *P* < 0.05; **, *P* < 0.01; ***, *P* < 0.001; ****, *P* < 0.0001.

Expanded ligand-receptor analysis demonstrated a profound increase in neutrophil-driven inflammatory communications in the MASLD liver. Elevated neutrophil ligands, such as *Thbs1*, *Mif*, *Ccl4*, *Ccl6*, and *Ccl9*, were paired with their corresponding receptors, including *Cd44*, *Cd74*, and *Ccr1*, on monocytes/macrophages and Kupffer cells (**Fig. 7B**), indicating amplifications of immune-fibrotic signaling pathways.

Consistent with these findings, gene expression analysis revealed that IF significantly decreased the expression of neutrophil-derived inflammatory cytokines and their receptors in control ASO-treated male livers, but not in *Hmgcs2* ASO-treated male livers (**Fig. 7C**). In females, IF failed to reduce these genes in either control ASO- or *Hmgcs2* ASO-treated female livers. Notably, *Hmgcs2* knockdown alone significantly elevated inflammatory cytokine transcript levels, consistent with fibrosis and neutrophil markers (**Fig. 5F** and **6F**), emphasizing a sex-dependent vulnerability in hepatic immune-fibrotic communications. Collectively, these findings demonstrate that IF-induced ketogenesis dampens neutrophil-mediated inflammatory pathways, thereby contributing to reduced hepatic fibrosis in males, whereas this protective communication axis is substantially blunted in females.

## DISCUSSION

Intermittent fasting (IF) has emerged as an effective dietary intervention with broad benefits against metabolic, cardiovascular, and neurological disorders (1). While its systemic effects have been well characterized (5-7; 16; 17), the underlying mechanisms remain incompletely defined. Recent human trials have begun to evaluate IF interventions (48–50); however, mechanistic insights continue to rely largely on rodent studies, especially those using male mice (4). Here we show that the metabolic benefits of IF are sexually dimorphic and partly dependent on hepatic ketogenesis. Although IF reduced obesity in both sexes, the effect was noticeably weaker in females. Importantly, these anti-obesity benefits occurred independently of hepatic ketogenesis in both males and females. In contrast, in male mice, IF ameliorated hepatic steatosis and fibrosis, and these effects were attenuated when ketogenesis was impaired. In female mice, IF conferred limited hepatic protection, and ketogenic insufficiency further increased susceptibility to metabolic stress.

Consistent with our findings, previous work has shown that females often exhibit reduced responsiveness to dietary interventions compared to males. For instance, long-term alternate-day fasting increases hepatic lipid accumulation in female mice but not males (51), while 9 hours of time-restricted feeding failed to prevent body-weight gain in females (52). Similarly, in humans, a very-low-carbohydrate ketogenic diet produced greater anti-obesity effects in men than in women (53), suggesting sex-dependent differences in nutrient utilization and energy metabolism in both humans and mice.

Physiological dimorphism during energy deprivation likely contributes to these outcomes (54; 55). We observed higher circulating ketone bodies in fasted females despite comparable hepatic expression of ketogenic enzymes, indicating sex differences in ketone production, utilization, or clearance rather than enzyme abundance. Prior work suggests that male livers preferentially oxidize lipids during fasting, whereas female livers maintain lipogenesis at the expense of amino acids (56). These features could blunt IF’s anti-steatotic action in females by sustaining lipid synthesis across fasting-feeding cycles. Moreover, because females generally exhibit less body-weight gain and milder hepatic steatosis under HFD feeding compared to males, their physiological window for metabolic adaptation during IF may be narrower, making improvements more difficult to detect. It will therefore be interesting in future studies to extend the duration of IF in female mice to determine whether longer interventions could reveal delayed or distinct metabolic responses.

A second key finding is that ketogenesis contributes differently to IF’s efficacy in males and females. Although we previously observed no sex differences in the spontaneous fatty liver phenotype of whole-body *Hmgcs2* knockout postnatal mice (21), our results suggest that adult female mice are more dependent on ketone body metabolism, which may be mediated through hepatic estrogen signalling, as previously suggested (57). In females, hepatic *Hmgcs2* knockdown exacerbated fibrosis and eliminated the benefit of IF, consistent with a greater dependence on ketone metabolism for hepatic homeostasis. In males, the anti-steatotic effects of IF were diminished but not abolished by loss of hepatic ketogenesis, indicating partial compensation by metabolic adaptations in extrahepatic tissues (5; 7; 12; 58; 59). Further studies are required to investigate sex differences in ketone body synthesis, utilization, and dependency upon dietary interventions.

Our mouse and human data together support a ketogenesis-inflammation-fibrosis axis. Co-expression analysis in the human liver revealed a positive association of *HMGCS2* with oxidative and peroxisomal metabolic pathways and a negative association with inflammatory, neutrophil, and fibrotic gene programs. In mice, single-cell and ligand-receptor analyses identified neutrophils as major senders of inflammatory signals to Kupffer cells and monocytes/macrophages. Both neutrophils and fibroblasts display distinct relationships with ketone bodies in both mouse and human systems. Neutrophils, being highly glycolytic and short-lived, do not depend on ketone oxidation for energy production but rather respond to extracellular ketone bodies as signaling molecules (60). Acetoacetate modulates inflammatory activity through the FFA2R receptor (61), whereas β-OHB has been shown to inhibit NLRP3 inflammasome activation (62), NETosis (60) and reactive oxygen species (ROS) production (63; 64), suggesting a non-metabolic immunoregulatory role of ketone bodies in neutrophil-mediated inflammation. Fibroblasts, by contrast, express enzymes required for ketone body metabolism and can modulate mitochondrial oxidative capacity in response to substrate availability, particularly under fasting or ketogenic conditions (65; 66). Consistent with these observations, our results showed that IF reduced neutrophil-derived inflammatory signaling in the male mouse liver, an effect that required intact hepatic ketogenesis. In contrast, female mouse livers exhibited limited suppression of inflammatory pathways by IF and exhibited increased cytokine expression when ketogenesis was impaired. Together with a previous study demonstrating that Kupffer cells utilize ketone bodies and that defective ketone oxidation in macrophages aggravates liver inflammation/fibrosis (26), our findings suggest that ketone bodies act as immunometabolic mediators in neutrophil-macrophage-stellate communication within the liver. Furthermore, sex-dependent variation in ketone body metabolism may underlie sex differences in immune responses, especially those mediated by macrophages, in the context of obesity-related nutritional stress or inflammation (67; 68).

In summary, our results demonstrate that hepatic ketogenesis serves as a key link between fasting metabolism and hepatic inflammation and fibrosis, and that its contribution to the benefits of IF is sex-dependent. These insights are critical for guiding the design of dietary and pharmacologic strategies that harness ketone pathways. Given that sex, age, and metabolic status can modulate the efficacy of dietary interventions, our findings, together with those of others, suggest that achieving optimal weight loss and metabolic health may require tailoring dietary strategies to specific target populations.

## ACKNOWLEDGEMENTS

This study was supported by the Diabetes Canada Award (OG-3-22-5697-KK) and partly by the Canadian Institutes of Health Research (CIHR) Project Grant (PJT 195676) to Dr. Kyoung-Han Kim. He is also a recipient of the National New Investigator Award from the Heart and Stroke Foundation of Canada (HSFC) and Early Researcher Award (ER22-17-236) from the Government of Ontario, Canada. The infrastructure was supported by the Canadian Foundation for Innovation, John R. Evans Leaders Fund (CFI-JELF; 37735) to Dr. Kyoung-Han Kim.

Termeh Aslani was supported by the University of Ottawa Heart Institute Endowed Scholarship. Shaza Asif was supported by the Canada Graduate Scholarships Master Award (CGS-M) by the Canadian Institutes of Health Research, and the Queen Elizabeth II Graduate Scholarship in Science and Technology. Yena Oh and Saif Dababneh were supported by the Frederick Banting and Charles Best Canada Graduate Scholarships Doctoral Awards (CGS-D). Cole Stocker was supported by the TRIANGLE CONNECT Summer Studentship. Dr. Sora Kwon was supported by the MITACS Elevate Fellowship (IT34864). Dr. Ri Youn Kim was supported by fellowships from the University of Ottawa Cardiology Research Endowment Fund and the National Research Foundation of Korea (NRF) by the Ministry of Education, Korea (2020R1A6A3A03039905). Dr. Joe Eun Son was awarded a Basic Science Research Program through the National Research Foundation of Korea, the Ministry of Education (2018R1A6A3A03012237).

## CONFLICT OF INTERESTS

The authors declare no conflicts of interest.

## AUTHOR CONTRIBUTIONS

T.A., S.A., R.Y.K., and K-H.K, conceived and designed research; T.A., S.A., H-J.L., S.K., S.D., R.Y.K., J.E.S., and K-H.K. performed experiments and analyzed data; Y.O., C.S., S.D., and K-H.K performed the bioinformatic analysis; T.A., S.A., Y.O., C.S., S.D., and K-H.K. prepared figures; J.P., X.Z. and J.E.S. provided technical support including histological analysis; A.E.M. provided reagents; M.F. and E.E.M. supported research funding acquisition and experimental design; L.T., provided research input/feedback as a patient partner; T.A., S.A., Y.O., C.S. and K-H.K. drafted and edited the manuscript. T.A., S.A., J.P., E.E.M. J.E.S. M.D.F. and K-H.K revised the manuscript; T.A., S.A., Y.O., H-J.L., C.S., S.K., J.P., S.D., X.Z., L.T., A.E.M., G.F.T., R.Y.K., M.D.F., E.E.M., J.E.S., and K-H.K. approved the final version of the manuscript.

## ABBREVIATIONS

AL: ad libitum
ASO: AntiSense Oligonucleotide BAT: Brown Adipose Tissue
β-OHB: β-hydroxybutyrate BW: Body Weight
H&E: Hematoxylin & Eosin HFD: High-Fat Diet
HMGCS2: 3-hydroxymethylglutaryl-CoA synthase 2 IF: Intermittent Fasting
IWAT: Inguinal White Adipose Tissues
PWAT: Perigonadal White Adipose Tissue
TG: Triglyceride
MASLD: Metabolic dysfunction-Associated Steatotic Liver Disease
scRNA-seq: single-cell RNA-sequencing
MASH: Metabolic dysfunction-Associated SteatoHepatitis
GTEx: Genotype-Tissue Expression

---

**Supplementary Figure 1.** Correlation analysis of hepatic HMGCS2 expression with metabolic and inflammatory genes in the human liver. Scatter plots showing correlations between hepatic *HMGCS2* expression and representative metabolic and inflammatory genes identified in Figure 7 from the GTEx human liver RNA-seq dataset (n = 226). Data points represent male (blue) and female (red) liver samples; black lines denote Spearman correlation trends with corresponding *r* and *P* values shown.

**Supplementary Figure 2.** Expression of hepatic ketogenic genes during fasting is not different between males and females. **(A)** Hepatic mRNA expression of *Acat1*, *Hmgcl*, and *Bdh1* in male and female C57BL/6J mice under fed or 24-hour fasted conditions. **(B)** Measurements of body weight in C57BL/6J male (left) and female (right) mice subjected to IF or AL for 3 weeks. Statistical analysis was performed by two-way ANOVA. **, *P* < 0.01.

**Supplementary Figure 3.** Antisense oligonucleotides (ASO) against *Hmgcs2* block fasting-induced ketogenesis with mRNA and protein knockdown. (A-B) Hmgcs2 gene (A) and protein (B) expression were tested by quantitative PCR and western blot. (C) Plasma β-OHB levels measured in C57BL6/J male mice in fed, fasted, fasted with bi-weekly ASO injection and fasted with weekly ASO injection (n = 3 / group). Statistical analysis was performed by one-way ANOVA. ***, *P* < 0.001; ****, *P* < 0.0001.

**Supplementary Figure 4.** Intermittent fasting and hepatic *Hmgcs2* knockdown do not affect cumulative food intake in male or female mice. Cumulative food intake (kcal per 3 days) measured over 20 weeks in male (A) and female (B) C57BL/6J mice treated with control or *Hmgcs2* ASO under *ad libitum* (AL) or intermittent fasting (IF) conditions.

**Supplementary Figure 5.** Intermittent fasting reduces adipose tissue mass and adipocyte size, with attenuated effects after *Hmgcs2* knockdown. (A-B) Weights of inguinal white adipose tissue (IWAT), perirenal white adipose tissue (PWAT), and brown adipose tissue (BAT) in male (A) and female (B) mice treated with control or *Hmgcs2* ASO under ad libitum (AL) or intermittent fasting (IF) conditions. (C-F) Representative H&E-stained IWAT sections and corresponding adipocyte size distributions in male (C, D) and female (E, F) mice. Insets show the mean adipocyte area. Scale bars, 100 µm. (G-H) *Lep* mRNA expression in male (G) and female (H) livers. Statistical analysis was performed by two-way ANOVA. *, *P* < 0.05; **, *P* < 0.01; ***, *P* < 0.001; ****, *P* < 0.0001.

**Supplementary Figure 6.** Cell-type identities in the mouse liver single-cell RNA-seq dataset. Expression of representative marker genes confirming the identities of major hepatic cell populations, including hepatocytes, hepatic progenitor cells, stellate cells, endothelial cells, Kupffer cells, and various immune cell subtypes. Marker gene expression was visualized across clusters to validate cell-type annotations used for downstream analyses.

## Notes

### Competing Interest Statement

The authors have declared no competing interest.

