## Supplementary figures and images for "Sex- and ketogenesis-dependent effects of intermittent fasting against diet-induced obesity and fatty liver disease"

### Supplementary Figure 1

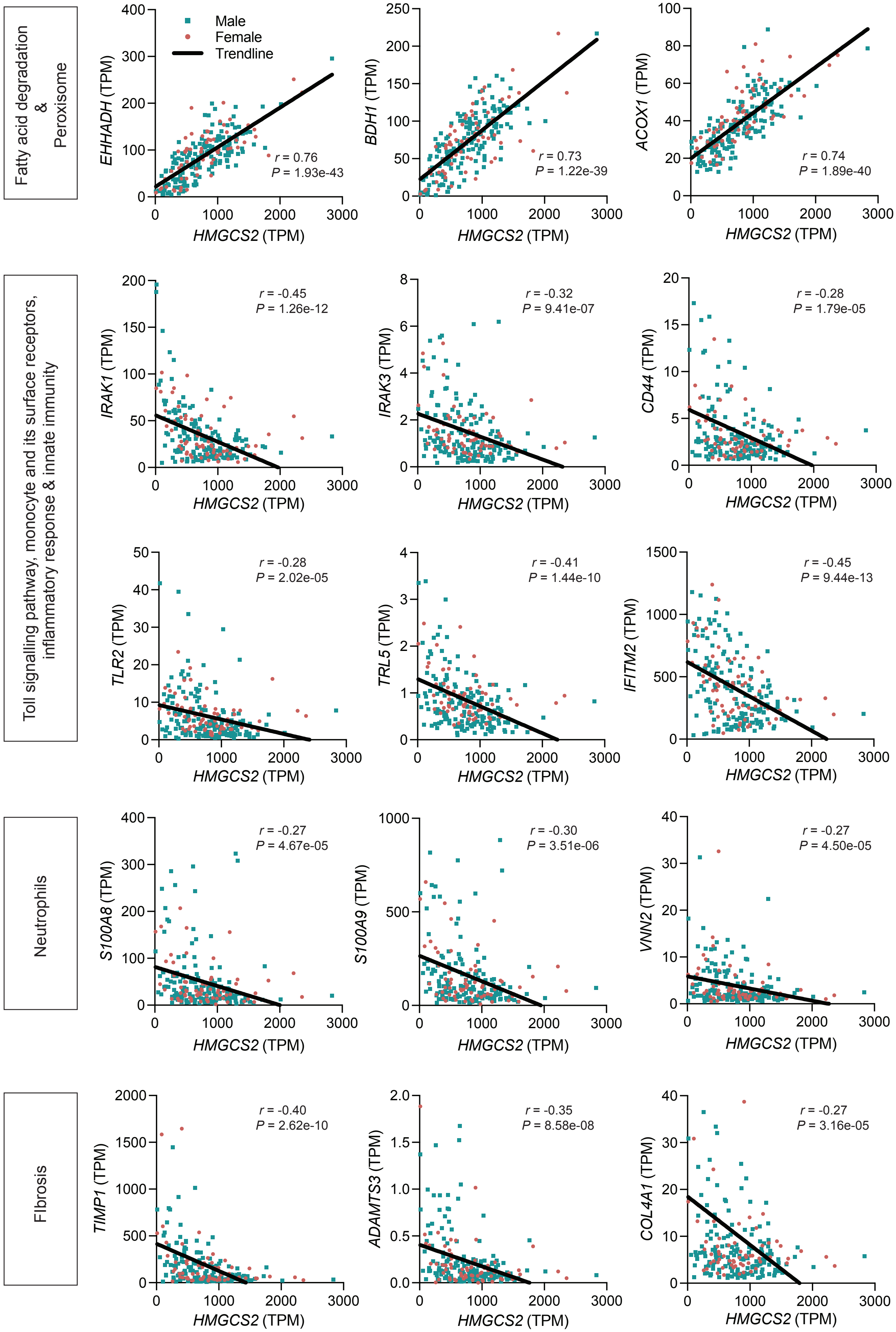

### Supplementary Figure 2

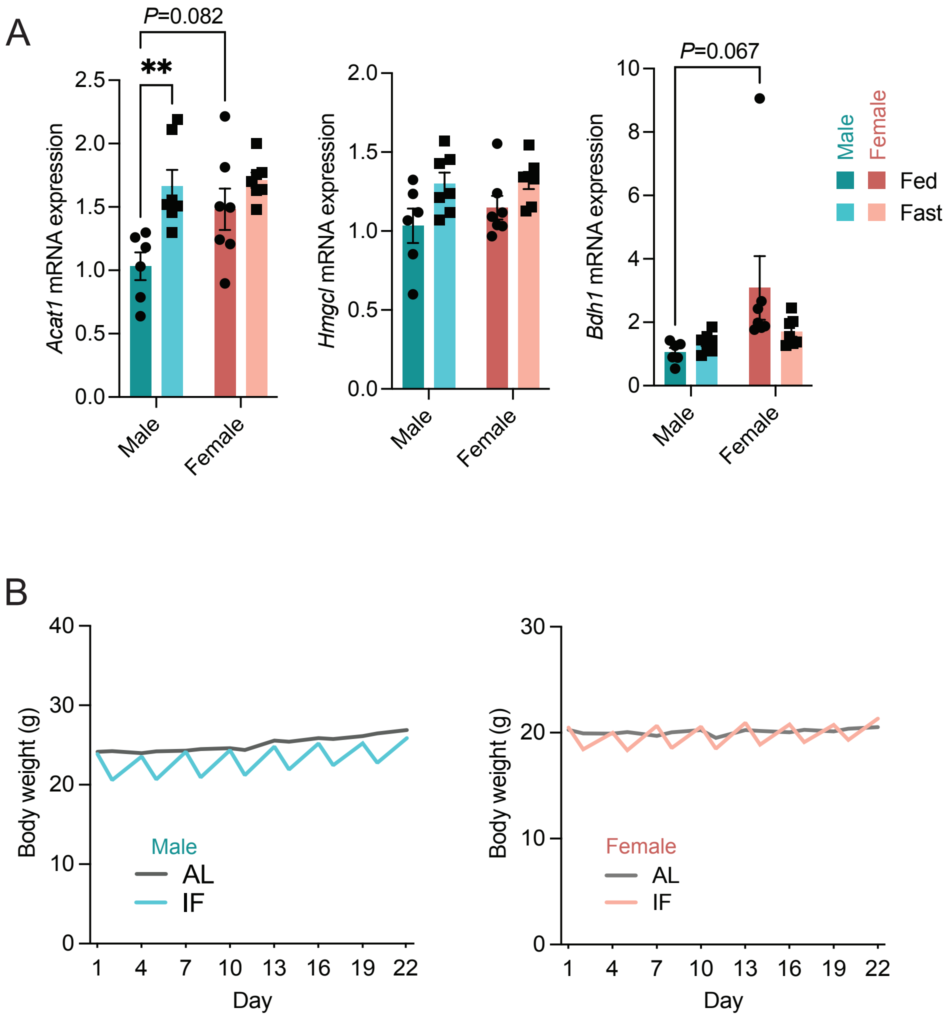

### Supplementary Figure 3

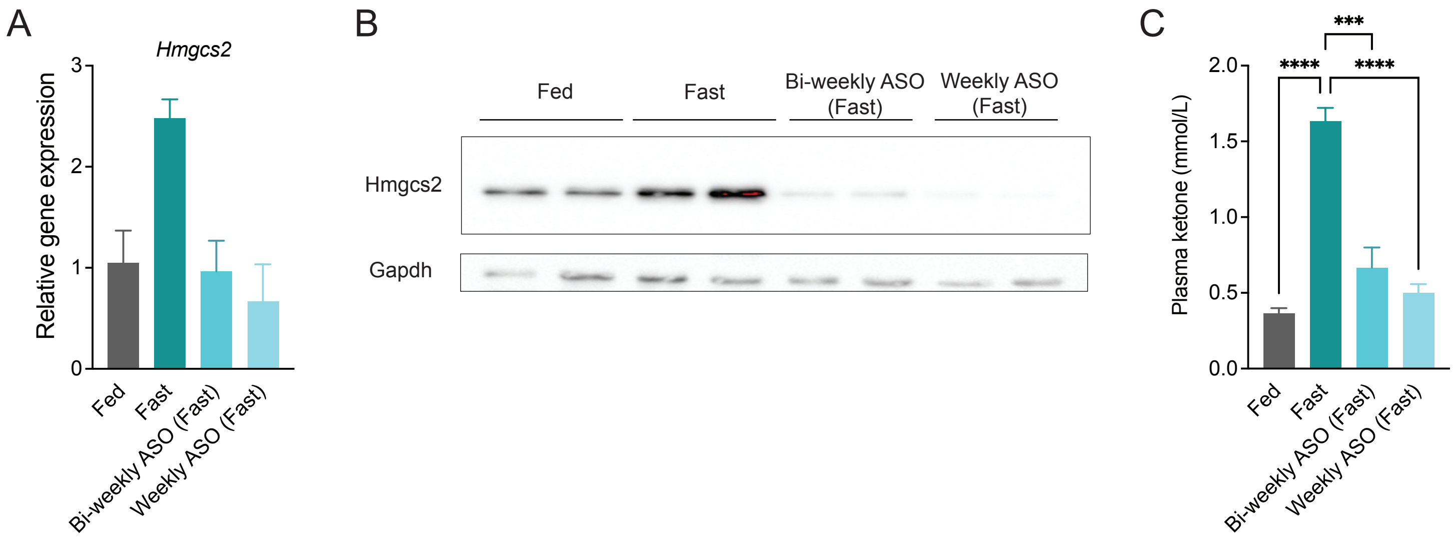

### Supplementary Figure 4

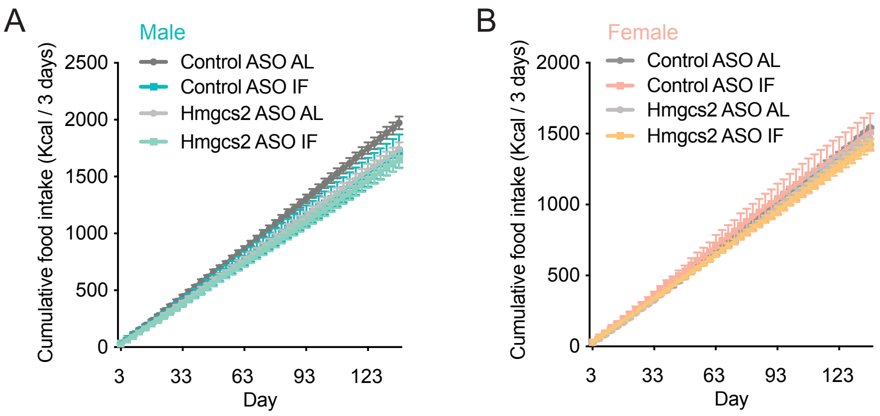

### Supplementary Figure 5

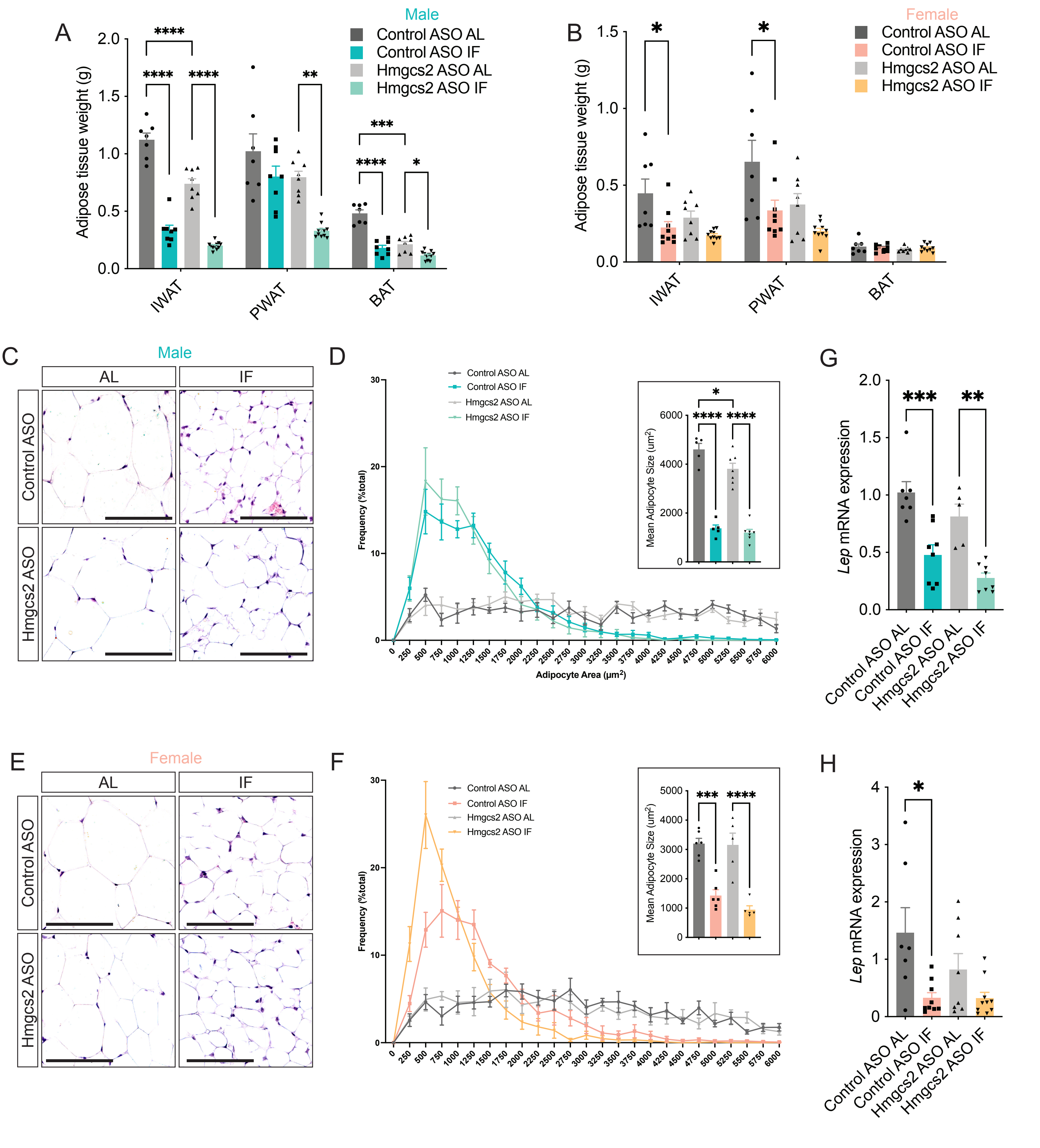

### Supplementary Figure 6

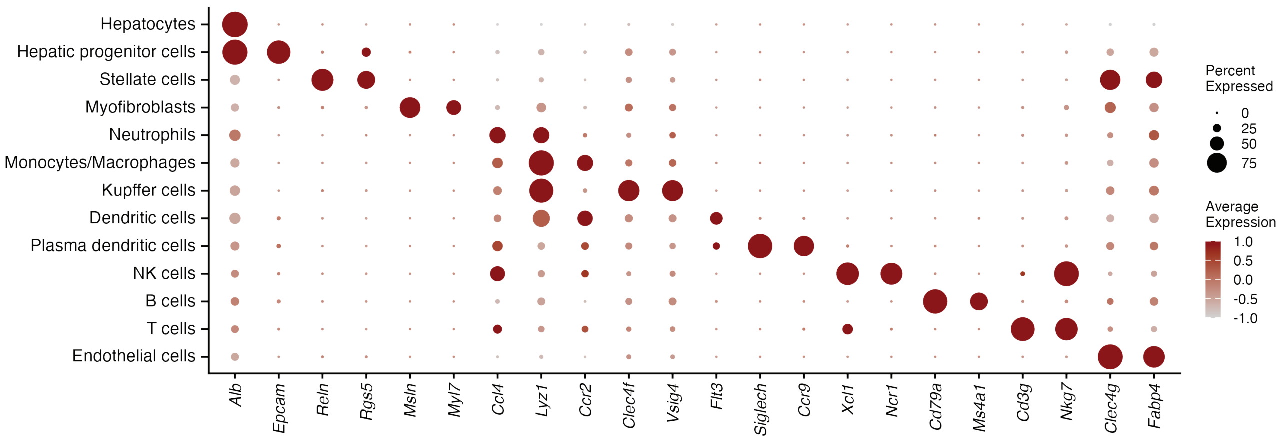
