## Supplementary Table 1 for "Sex- and ketogenesis-dependent effects of intermittent fasting against diet-induced obesity and fatty liver disease"

Supplementary Table 1. List of qPCR primers

| Gene | Forward (5' – 3') | Reverse (5' – 3') |
|---|---|---|
| *Hmgcs2* | GGTGTCCCGTCTAATGGAGA | ACACCCAGGATTCACAGAGG |
| *Bdh1* | GAATTCAGCCTGCCGGTTTG | TGCATCCCGCTGTCAGGTAA |
| *Lep* | GATGGACCAGACTCTGGCAG | AGAGTGAGGCTTCCAGGACG |
| *Pparg* | CACCAGTGTGAATTACAGCAAATC | AGCTGATTCCGAAGTTGGTG |
| *Plin2* | GACCTTGTGTCCTCCGCTTAT | CAACCGCAATTTGTGGCTC |
| *Fsp27* | CTGGAGGAAGATGGCACAAT | GGGCCACATCGATCTTCTTA |
| *Col1a1* | ACGTGGAAACCCGAGGTATG | TTGGGTCCCTCGACTCCTAC |
| *Timp1* | CGAGACCACCTTATACCAGCG | ATGACTGGGGTGTAGGCGTA |
| *Cd44* | ACTAGATCCCTCCGTTTCATCC | GGTTACATTCAAATCGATCTGCTG |
| *Tbp* | GAAGCTGCGGTACAATTCCAG | CCCCTTGTACCCTTCACCAAT |
| *Col1a2* | GTAACTTCGTGCCTAGCAACA | CCTTTGTCAGAATACTGAGCAGC |
| *S100a6* | TTCTCGTGGCCATCTTCCAC | GCCAATGGTGAGCTCCTTCT |
| *Rgs5* | GGGTTGCCTGTGAGAATTACAAG | CCTCTGTCTGGATGAATTCTTCATAA |
| *Pdgfa* | TGGCTCGAAGTCAGATCCACA | TTCTCGGGCACATGGTTAATG |
| *Ly6g* | TGTGCTCATCCTTCTTGTGG | AGGGGCAGGTAGTTGTGTTG |
| *S100a8* | CAAGGAAATCACCATGCCCTCTA | ACCATCGCAAGGAACTCCTCGA |
| *S100a9* | TGGTGGAAGCACAGTTGGCAAC | CAGCATCATACACTCCTCAAAGC |
| *Mmp8* | TGGCATTCAGACAATCTATGGACCT | CACGGAGTGTGGTAGTAGCATCAA |
| *Thbs1* | AGTGGAAGAGCATCACGCTG | CACCACGTTGTTGTCAAGGG |
| *Mif* | TGCCCAGAACCGCAACTACAGTAA | TCGCTACCGGTGGATAAACACAGA |
| *Ccl4* | CAAACCTAACCCCGAGCAACAC | GGTCTCATAGTAATCCATCACAAAGC |
| *Ccl6* | ATGAGAAACTCCAAGACTGCC | TTATTGGAGGGTTATAGCGACG |
| *Ccl9* | CCCTCTCCTTCCTCATTCTTACA | AGTCTTGAAAGCCCATGTGAA |
| *Cd74* | GCTGGATGAAGCAGTGGCTCTT | GATGTGGCTGACTTCTTCCTGG |
| *Ccr1* | ACTGCTGTAAGAGCCTTTGGG | AGCACCAGAATCACTAGGACA |
| *Acat1* | GTCTGGCTAGTATTTGCAAGG | TTCAGCCGGTCACATGG |
| *Hmgcl* | GGACTTCATCTGTCAAGCC | TCATTGTATACACCCAATTCCC |
